# From molecular lipidomics to interpretable food lipid profiles: the Lipid Food Profile module in LipidOne

**DOI:** 10.64898/2026.06.15.732299

**Authors:** Dorotea Frongia Mancini, Husam B.R. Alabed, Roberto Maria Pellegrino

## Abstract

LC/MS-based food lipidomics provides detailed information on intact lipid species, but the resulting datasets are often difficult to translate into concepts directly useful for food quality, processing, nutritional profiling and authenticity assessment. Here, we present Lipid Food Profile (LFP), a module of the LipidOne platform designed to convert annotated LC/MS lipidomics data into interpretable food-relevant lipid indices. LFP applies an in silico hydrolysis strategy to reconstruct acyl, alkyl and alkenyl chains from intact lipid species while preserving their lipid-class origin. The reconstructed information is then summarized into index categories related to food lipid quality, compositional balance, omega balance, oxidative stability, chain remodelling and ether-linked chain contribution.

The interpretative value of LFP was evaluated using three published food lipidomics datasets addressing different analytical questions: X-ray-induced lipid remodelling in *Chlorella vulgaris*, spatial lipid heterogeneity in *Mugil cephalus* bottarga, and geographical-origin assessment of camel milk. Across these case studies, LFP recovered the main conclusions of the original lipidomics investigations, including treatment-associated lipid remodelling, inner–outer layer differences in bottarga and regional variation in camel milk. Importantly, LFP reorganized these findings into a smaller number of food-oriented indices, providing additional information on saturation balance, oxidative susceptibility, chain architecture and classification potential.

Overall, LFP provides an interpretative layer for LC/MS food lipidomics that complement conventional fatty-acid analysis and molecular-species-based interpretation. By translating complex lipidomic tables into structured lipid index profiles, the module may support more accessible and chemically meaningful analysis of food composition, processing effects, lipid quality and exploratory traceability applications. LFP is freely accessible through the LipidOne web platform (LipidOne.eu).

**Highlights:**

- Lipid Food Profile translates LC/MS food lipidomics into interpretable lipid indices.
- The workflow preserves chain and lipid-class information without chemical hydrolysis.
- Published case studies show that LFP recovers and extends previous interpretations.
- LFP supports food quality, processing and exploratory origin/authenticity assessment.
- The module complements conventional fatty-acid analysis and molecular lipidomics.

## 1. Introduction

Lipids are major determinants of food composition, nutritional value, sensory properties, technological performance and oxidative stability. In edible matrices, the lipid fraction contributes to energy density, palatability, texture, shelf life, processing behaviour, product differentiation and authenticity. For this reason, lipid characterization is central to food science, including quality control, comparison of food matrices, evaluation of processing effects, assessment of geographical or botanical origin and nutritional profiling. Recent advances in food lipidomics have expanded this field by enabling detailed analysis of lipid classes and molecular species across foods and food-derived products (Muguruma et al., 2022; Song et al., 2022; Tietel et al., 2023).

Traditionally, food lipids have been described mainly through fatty-acid analysis. Lipids are hydrolysed or transesterified, converted into fatty acid methyl esters and analysed by GC-FID or GC-MS. This robust and standardized approach supports the calculation of lipid-quality indices, including the atherogenic index, thrombogenic index, hypocholesterolaemic/hypercholesterolaemic ratio, PUFA/SFA ratio, unsaturation index and peroxidability index. These variables condense fatty-acid profiles into interpretable metrics related to lipid quality, compositional balance and oxidative susceptibility (Chen & Liu, 2020; Ulbricht & Southgate, 1991).

However, conventional fatty-acid profiling removes the molecular and lipid-class context of the original lipidome. Chains originating from triacylglycerols, phospholipids, cholesteryl esters, sphingolipids or ether-linked lipids are collapsed into a single pool, preventing the user from determining the lipid classes in which specific chains were originally embedded. Moreover, chemical conversion is not equivalent across all lipid classes, and ether-linked lipids, plasmalogens and sphingolipids may require specific analytical considerations (Delmonte et al., 2020; Gómez-Cortés et al., 2019; Ichihara & Fukubayashi, 2010; Morrison & Smith, 1964).

Liquid chromatography-mass spectrometry has therefore become increasingly important in food lipidomics because it preserves lipid information and enables the detection of molecular species across multiple lipid classes. LC/MS-based lipidomics can reveal class-specific patterns associated with origin, processing, storage, oxidation or formulation. At the same time, these datasets often consist of long lists of lipid species that are chemically informative but not immediately readable using concepts commonly applied in food chemistry.

To address this gap, we developed Lipid Food Profile (LFP), a new module of the LipidOne web platform (Alabed et al., 2024, 2026; Pellegrino et al., 2022) designed to translate LC/MS food lipidomics datasets into food-relevant lipid indices. LFP applies a class-aware in silico hydrolysis strategy in which annotated intact lipid species are computationally resolved into their constituent acyl, alkyl and alkenyl chains while preserving lipid-class origin. The reconstructed chain-level information is then converted into 20 indices grouped into six categories related to food lipid quality, chain remodelling, compositional balance, oxidative stability, omega balance and ether-linked chain contribution. Rather than replacing conventional fatty-acid analysis, LFP adds an interpretative layer to intact-lipid datasets, allowing food scientists to move from complex molecular-species tables to compact and chemically meaningful food lipid profiles.

## 2. Translating LC/MS lipidomics into food lipid profiles

Lipid Food Profile was developed for LC/MS lipidomics datasets annotated at the molecular-species level. The module starts from a lipid abundance matrix in which food samples are represented by annotated lipid species expressed in standard shorthand nomenclature, because LFP requires sufficient structural information to resolve each lipid species into its constituent chains. Datasets limited to lipid-class abundances or unresolved sum compositions do not retain enough information for complete chain reconstruction.

The first step is the computational reconstruction of lipid building blocks. Each annotated lipid is decomposed into its acyl, alkyl or alkenyl chains while retaining the lipid class from which each chain originates. This representation allows food lipidomes to be examined both as global chain pools and as class-resolved lipid structures. We refer to this procedure as in silico hydrolysis because it generates a chain-level representation of the lipidome without chemical hydrolysis. Chemical hydrolysis or transesterification followed by GC-based analysis measures experimentally released fatty-acid derivatives and remains essential for standardized quantitative fatty-acid profiling. In contrast, in silico hydrolysis infers chain distributions from LC/MS annotations and depends on lipid coverage, annotation quality and the semi-quantitative comparability of the original dataset. Its main advantage is the preservation of molecular and lipid-class context, including ether-linked chains when present, which are not readily resolved by conventional chemical hydrolysis.

After chain reconstruction, LFP converts chain-level information into curated food-relevant indices. Classical lipid-quality indices are retained because they are widely used in food-fat evaluation; however, in LFP, they are recalculated from LC/MS-derived chains and interpreted as compositional indices of the analysed food lipidome rather than as direct physiological predictors. These variables can then be compared across food samples, origins, processing conditions or treatments using statistical approaches commonly applied to lipid species, but with greater interpretative compactness.

A distinctive feature of the LFP workflow is the connection between numerical values of the indices and concise interpretation phrases. Each index is associated with predefined foodomics-oriented statements describing the meaning of increased or decreased values relative to a reference condition. These statements are not intended to generate deterministic health claims, but to translate statistical variation into chemically meaningful language, such as increased unsaturation, altered omega balance, higher oxidative susceptibility, modified chain-length distribution or greater contribution of ether-linked structures.

As illustrated in Figure 1, lipidomics input data are processed within the LFP module through lipid parsing and feature extraction, followed by the calculation of 20 LFP indices grouped into six index categories. These indices are then used as variables for statistical analysis, ultimately supporting lipid food characterization.

**Figure 1:**
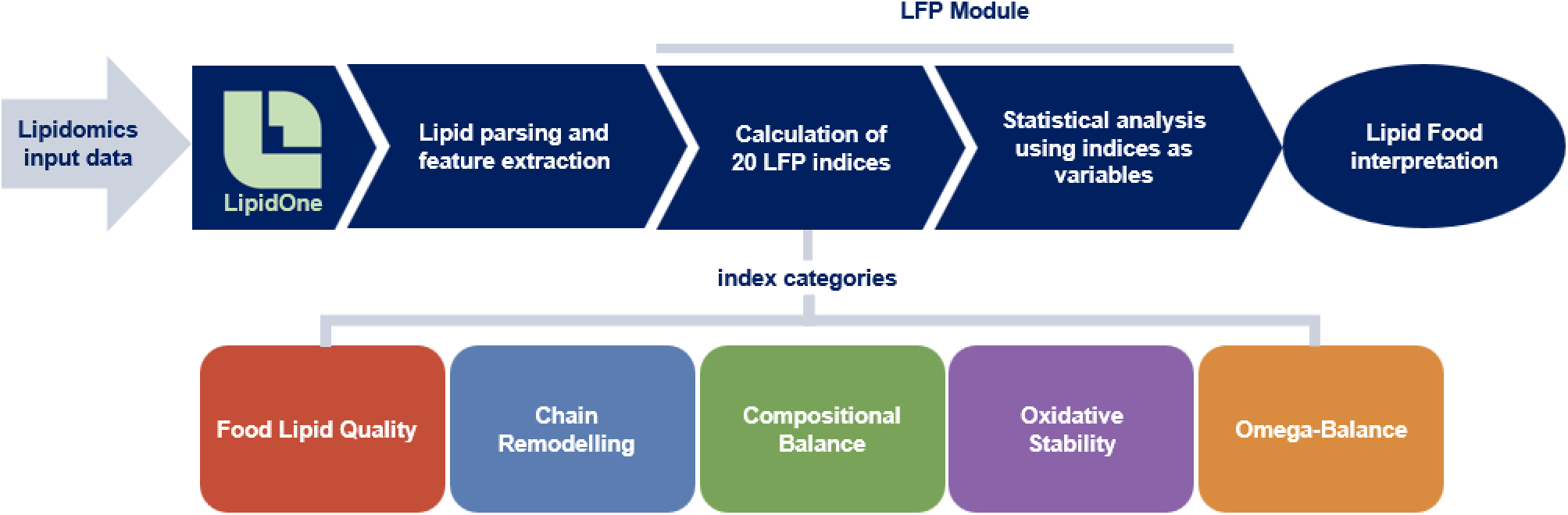
Schematic overview of the Lipid Food Profile workflow. Annotated LC/MS lipidomics data are processed through lipid parsing, class-aware in silico hydrolysis and reconstruction of acyl, alkyl and alkenyl chains. The resulting information is converted into 20 food-relevant indices grouped into six interpretative categories and then used for statistical analysis and food lipid characterization.

## 3. LFP index categories and foodomics interpretation

After computational chain reconstruction, LFP condenses LC/MS lipidomics datasets into a compact set of chemically interpretable indices. The index panel was assembled by adapting established fatty-acid quality metrics and lipid chemistry indices to chain-level data reconstructed from intact LC/MS annotations. A key feature of LFP is that indices can be calculated globally, across the entire lipidome, or within specific lipid classes when the underlying data support such analysis.

For practical interpretation, the indices were grouped into six foodomics-oriented categories: Food Lipid Quality, Compositional Balance, Omega Balance, Oxidative Stability, Chain Remodelling and Ether-Linked Chains (Table 1). These categories are not intended to represent rigid biochemical compartments, but rather to organize indices according to their dominant meaning in food lipid analysis. The ether-linked category represents an extension enabled by LFP, because in silico hydrolysis reconstructs not only acyl chains but also alkyl and alkenyl moieties. This allows ether-linked chains to be considered as compositional features potentially relevant to lipid organization, matrix specificity and oxidative behaviour, without implying direct nutritional claims.

**Table 1.** Foodomics-oriented index categories implemented in Lipid Food Profile, with their main interpretative meaning, corresponding LFP indices, and supporting literature references.

| Index category | Main foodomics-oriented interpretation | LFP index | Literature reference(s) |
| --- | --- | --- | --- |
| **Food Lipid Quality** | General lipid-quality balance derived from reconstructed chain composition. | AI | Chen & Liu, 2020; Dal Bosco et al., 2024 |
|  |  | TI | Dal Bosco et al., 2024; Paszczyk & Czarnowska-Kujawska, 2022; Rashidimehr et al., 2025 |
|  |  | h/H ratio | Chen & Liu, 2020; Santos-Silva et al., 2002 |
| **Compositional Balance** | Overall distribution of saturation level and chain architecture. | Long-Chain FA % | Paszczyk et al., 2019 |
|  |  | MCFA/LCFA | Sun et al., 2013 |
|  |  | Medium-Chain FA % | Sun et al., 2013 |
|  |  | MUFA/SFA | Yazdanparast et al., 2025 |
|  |  | PUFA/MUFA | Hamdy et al., 2026 |
|  |  | PUFA/SFA | Chen & Liu, 2020 |
|  |  | Saturated FA % | Kaçar et al., 2024 |
| **Omega Balance** | Balance among nutritionally relevant unsaturated-chain families. | DHA/ARA | Araújo et al., 2022 |
|  |  | EPA/ARA ratio | Araújo et al., 2022 |
|  |  | EPA+DHA fraction | Chen & Liu, 2020; Nava et al., 2023; Sprague et al., 2016 |
| **Oxidative Stability** | Susceptibility of the lipidome to oxidation based on unsaturation pattern. | Peroxidability index | Yun & Surh, 2012 |
|  |  | Unsaturation index | Chen & Liu, 2020 |
| Chain Remodelling | Changes in reconstructed chain architecture across samples or treatments. | C16 Desaturation Index | Inostroza et al., 2022 |
|  |  | C18 Desaturation Index | Inostroza et al., 2022 |
| Ether-Linked Chains | Contribution of ether-linked moieties retained by intact-lipid annotation. | Ether Chain Fraction | Z. Chen et al., 2025 |
|  |  | Alkyl-chain Fraction | Z. Chen et al., 2025 |
|  |  | Alkenyl-chain Fraction | Z. Chen et al., 2025 |

By transforming reconstructed chains into these index categories, LFP compresses complex intact-lipid datasets into a chemically interpretable profile that can be used in univariate, multivariate, clustering and classification analyses. The complete list of indices, formulas, interpretative phrases and bibliographic references are reported in Supplementary Table S1.

### 3.1. Analytical functions available in the LFP module

The LFP module was designed to provide users with a set of complementary analytical tools for the exploration, interpretation and classification of food lipidomic profiles. Rather than focusing exclusively on individual lipid species, the module translates the lipidome into a reduced set of food-relevant indices, allowing the user to investigate compositional differences from a more interpretable perspective. The available functions (Table 2) include univariate statistical tools, multivariate analysis, clustering approaches and classification utilities, which together support different analytical needs, from the comparison of experimental groups to the evaluation of food origin or authenticity.

**Table 2.** Overview of the analytical functions available in the LFP module. The table summarizes the main groups of functions provided by the module and their intended use for the exploration, interpretation and classification of food lipidomic profiles.

| Group of analysis functions | Available functions | Analytical purpose |
| --- | --- | --- |
| **Univariate statistical analysis** | Bar graph; Summary; Volcano plot; Bubble plot; Category bar plot; Trajectory plot | Comparison of individual indices or lipid features among experimental groups; visualization of significant changes, trends and category-level patterns. |
| **Multivariate statistical analysis** | PCA; PLS-DA | Exploration of sample distribution, group separation and main variables contributing to lipid profile differences. |
| **Clustering analysis** | Heatmap | Visualization of coordinated LFP index patterns across samples or groups, supporting the recognition of similarities and differences in food lipid profiles. |
| **Food origin and authenticity assessment** | Classification performance; Unknown samples classifier | Evaluation of supervised classification models and assignment of unknown samples to predefined food groups or origins. |

**Univariate functions**, such as bar graphs, volcano plots, bubble plots and category-level summaries, allow users to identify indices or lipid features that differ significantly between groups and to visualize their biological or technological relevance in an intuitive way.

**Multivariate tools**, including PCA and PLS-DA, provide an overview of sample distribution and group separation, helping to assess whether the lipid functional profile captures systematic differences among food samples. **Clustering** visualizations further support the recognition of patterns within the dataset, highlighting similarities among samples or groups without requiring a priori assumptions.

In addition, the LFP module includes classification-oriented functions specifically intended for food **origin and authenticity assessment**. These tools allow users to evaluate the performance of supervised models and to classify unknown samples according to reference lipidomic profiles. In this context, the module can be used not only to explore differences among known food groups, but also to support the assignment of new samples to predefined categories.

Overall, the LFP module provides a flexible analytical workflow that combines statistical comparison, visual interpretation and classification, making lipidomic data more accessible for food quality, origin, authenticity and nutritional studies.

## 4. Case-study evaluation of LFP interpretative value

To evaluate the interpretative value of LFP, we applied the module to three published LC/MS-based food lipidomics datasets representing different matrices and analytical questions. The first dataset concerns lipid remodelling in Chlorella vulgaris exposed to increasing X-ray doses (Casula et al., 2024); the second describes spatial lipid heterogeneity in Mugil cephalus bottarga (Manis et al., 2026); and the third reports geographical variation in camel milk from three ecologically distinct regions (Zhu et al., 2026). These datasets were selected because the original studies reported complex lipidomic changes at the molecular-species or lipid-class level, while addressing questions relevant to food quality, processing, compositional structure or origin assessment.

The objective was to assess whether index-based interpretation could recover the principal conclusions of the original studies and reorganize them into a compact foodomics framework.

### 4.1. Chlorella vulgaris irradiation dataset

As a first case study, we re-analysed the lipidomics dataset published by Casula et al. on Chlorella vulgaris exposed to increasing X-ray doses. In the original study, irradiation did not substantially impair growth, but induced marked lipid remodelling involving fatty acids, thylakoid lipids, phospholipids and triacylglycerols. These changes were interpreted as part of an adaptive response to radiation stress, characterized by alterations in saturation level, membrane lipid composition and storage lipid accumulation.

LFP recovered the main trends described in the original work, while reorganizing them into a reduced set of index domains. At the highest interpretative level, the category-level trajectory summary showed that the lipidomic response to irradiation was not uniformly distributed across all index families (Figure 2A). Chain Remodelling displayed the strongest negative trajectory across irradiated samples, whereas Oxidative Stability and Compositional Balance also decreased relative to the control. In contrast, Food Lipid Quality showed a positive trajectory, while Omega Balance and Ether-Linked Chains remained comparatively stable. This result indicates that irradiation mainly affected indices related to chain architecture, saturation balance and oxidative susceptibility, rather than producing a generalized shift across the whole LFP panel.

**Figure 2:**
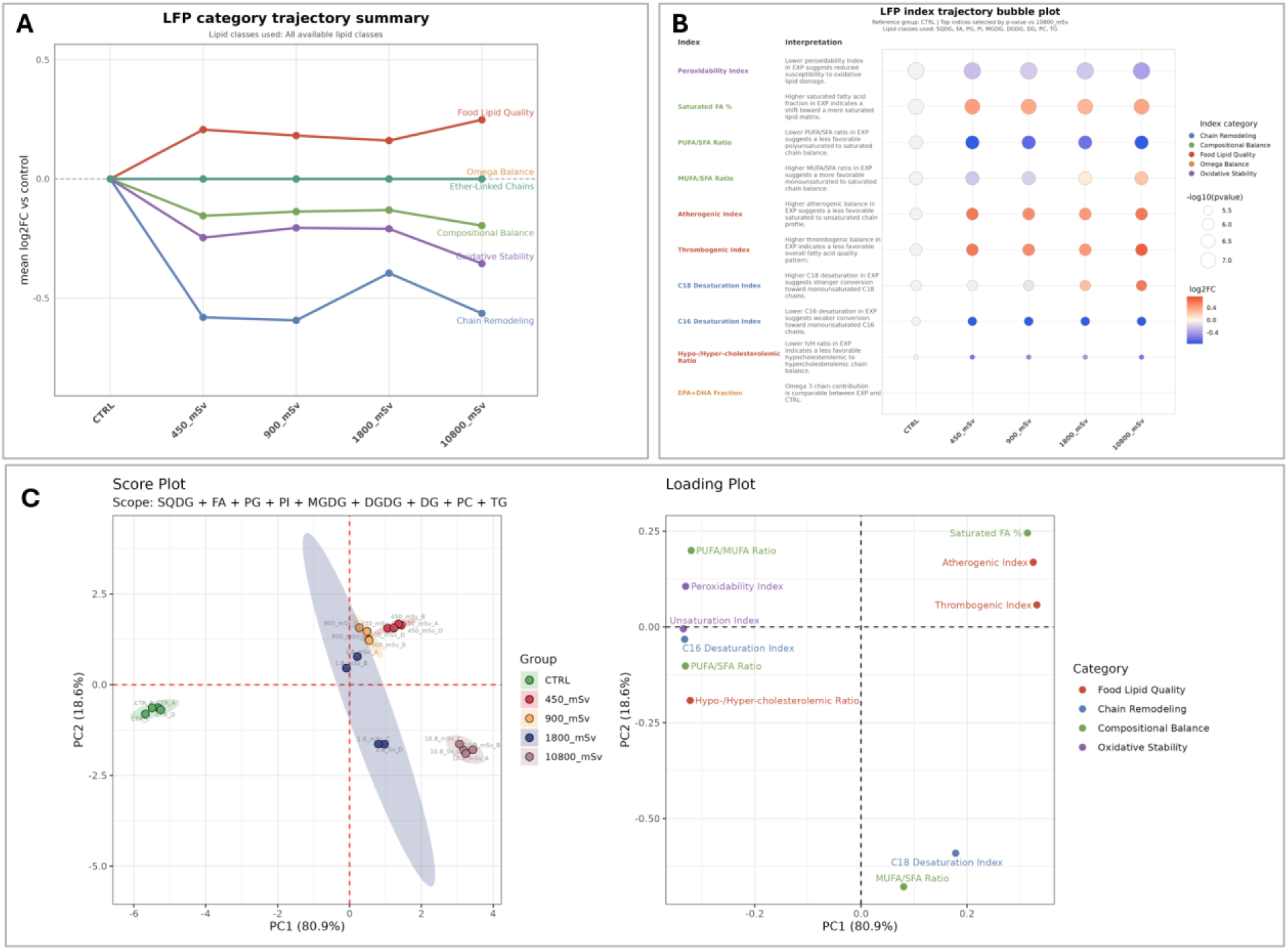
Chlorella vulgaris irradiation case study analysed using LFP indices. (A) Index-category trajectory summary showing treatment-related changes across LFP categories. (B) Index trajectory bubble plot highlighting the most responsive indices across irradiation doses. (C) PCA score and loading plots showing the separation of control and irradiated samples in the LFP index space and the indices contributing to this separation.

At the individual-index level, the bubble plot confirmed that the response was driven by a restricted subset of indices rather than by uniform changes in all calculated variables (Figure 2B). Indices related to the balance between saturated, monounsaturated and polyunsaturated chains, together with indices associated with oxidative susceptibility, were among the most responsive features. In particular, changes in the peroxidability index, saturated fatty-acid percentage, PUFA/SFA ratio, MUFA/SFA ratio and desaturation-related indices contributed to the interpretation of irradiation-associated lipid remodelling. These index-level patterns are consistent with the original observation that X-ray exposure modified the relative abundance of saturated, monounsaturated and polyunsaturated chains and affected lipid classes involved in both membrane structure and storage metabolism.

The multivariate score and loading plots further supported the ability of LFP indices to capture irradiation-associated lipid remodelling (Figure 2C). In the score plot, control samples clustered separately from irradiated samples, mainly along PC1, while the irradiated groups showed a dose-related redistribution within the LFP index space. The corresponding loading plot indicated that this separation was associated with opposing contributions of lipid-quality and compositional indices. Saturated fatty-acid percentage, atherogenic index and thrombogenic index contributed positively to PC1, whereas indices related to unsaturation and polyunsaturated-chain balance, including peroxidability index, unsaturation index, PUFA/SFA ratio and PUFA/MUFA ratio, contributed in the opposite direction. This pattern reinforces the interpretation that irradiation reshaped the reconstructed chain profile by modifying saturation balance, desaturation-related features and oxidative susceptibility.

The main advantage of this representation is that it makes radiation-induced remodelling easier to interpret at the foodomics level. Rather than requiring the user to inspect multiple lipid classes and individual molecular species separately, LFP summarizes the response as a coordinated shift involving saturation balance, chain remodelling and oxidative vulnerability. Thus, the Chlorella case study shows that LFP can reproduce the principal conclusions of the original lipidomics investigation while organizing them into an index-based framework directly related to food lipid composition and stability.

### 4.2. Bottarga spatial lipidomics dataset

As a second case study, we re-analysed the spatial LC–HRMS lipidomics dataset reported by Manis et al. on Mugil cephalus bottarga. The original study identified pronounced lipid heterogeneity between the inner and outer layers of the product. The inner region was characterized by higher levels of complex lipids, including cardiolipins, phosphatidylcholines, phosphatidylethanolamines and triacylglycerols, whereas the outer region showed greater abundance of lysophospholipids, free fatty acids and other lipid degradation products. This spatial pattern was interpreted as reflecting differential preservation, hydrolysis and oxidative exposure associated with salting and drying.

When the dataset was transformed into the LFP index space, the overall food lipid shift between the external and internal bottarga layers was not uniformly distributed across all index categories (Figure 3A). The strongest category-level differences were observed for Chain Remodelling and Compositional Balance, followed by Ether-Linked Chains and Oxidative Stability, whereas Food Lipid Quality and Omega Balance showed smaller shifts. This indicates that the spatial heterogeneity of bottarga was mainly captured by indices describing reconstructed chain architecture and compositional organization, rather than by a homogeneous change across the entire LFP panel.

**Figure 3:**
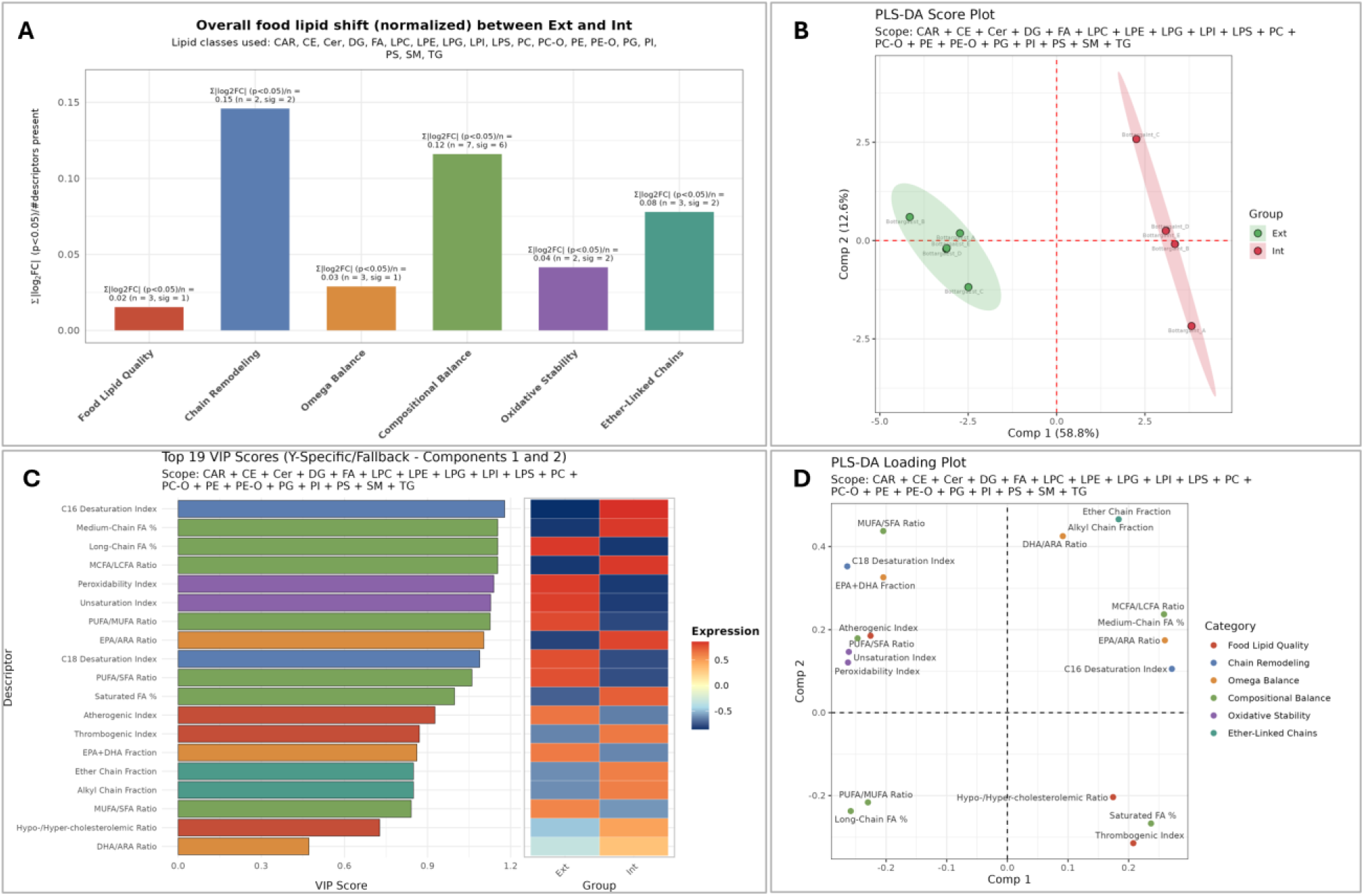
Bottarga spatial lipidomics case study analysed using LFP indices. (A) Overall food lipid shift between external and internal bottarga layers across LFP index categories. (B) PLS-DA score plot showing separation between external and internal layers in the LFP index space. (C) VIP score plot identifying the most discriminant LFP indices contributing to layer separation. (D) PLS-DA loading plot showing the direction and contribution of individual indices to the separation between external and internal layers.

The inner and outer bottarga layers also remained clearly distinguishable by PLS-DA, with the two regions separating mainly along the first component (Figure 3B). This indicates that the spatial divergence originally observed at the molecular-species level was retained after index compression and became interpretable in terms of a smaller number of food-relevant variables.

The VIP score plot confirmed that the most discriminant indices were concentrated in a restricted subset of the LFP panel rather than being uniformly distributed across all variables (Figure 3C). The highest-ranking indices included the C16 desaturation index, medium- and long-chain fatty-acid percentages, MCFA/LCFA ratio, peroxidability index, unsaturation index, PUFA/MUFA ratio, EPA/ARA ratio, C18 desaturation index and PUFA/SFA ratio. This pattern indicates that the spatial lipid heterogeneity of bottarga was mainly explained by chain remodelling, compositional balance and oxidative susceptibility, with additional selective contributions from Food Lipid Quality, Omega Balance and Ether-Linked Chains.

The loading structure further clarified the direction of the indices contributing to the separation between external and internal layers (Figure 3D). Indices related to C16 and C18 desaturation, saturation balance, medium-versus long-chain distribution, PUFA/SFA and MUFA/SFA ratios, and broader compositional remodelling contributed to the spatial separation. These findings support the idea that the inner-versus-outer gradient in bottarga can be understood not only as a list of altered lipid species, but also as a coordinated change in reconstructed chain architecture.

Overall, these findings are coherent with the original interpretation, in which the inner layer retained a lipid profile richer in complex lipids, whereas the outer layer displayed a more transformed signature associated with processing-related exposure. This hierarchical view represents one of the main added values of LFP: the module does not simply confirm that the two regions differ but identifies which index domains explain most of the difference. In practical terms, LFP compresses complex intact-lipid data into a compact set of interpretable categories, allowing spatial lipid heterogeneity to be described in terms of chain remodelling, compositional balance and oxidative stability.

### 4.3. Camel milk geographical origin assessment

As a third case study, we evaluated the Food Origin/Authenticity Assessment section of LFP using a UHPLC-MS/MS lipidomics dataset of Bactrian camel milk collected from three regions of Xinjiang, China: Fuhai County (AC), Yumin County (TC) and Huocheng County (LC). The original study reported regional differences involving several lipid classes, including glycerophospholipids, glycerolipids, sphingolipids and PUFA-derived oxidized lipids. Thus, this dataset was suitable to test whether LFP could preserve geographical information while translating molecular lipidomics data into a more interpretable index-based space.

A key advantage of LFP in this context is that in silico hydrolysis is class-aware. Reconstructed chains are not treated as an anonymous fatty-acid pool, but remain linked to the lipid classes from which they derive. This allows users to include or exclude lipid classes before calculating food-relevant indices, focusing the analysis on the lipid fraction that is most informative for the question being addressed. The lipid-class overview confirmed that regional variation was not uniformly distributed across the lipidome, with only a subset of classes showing significant differences among AC, TC and LC (Figure 5A). These informative classes were therefore retained for the main classification workflow, whereas the corresponding all-class analysis was reported as supplementary material.

Using the selected lipid-class subset, the PLS-DA classification map showed a clear organization of the samples according to geographical origin, with AC, TC and LC occupying distinct regions of the LFP index space (Figure 4A). Cross-validation supported the stability of this classification, with high accuracy and balanced accuracy across the selected model configuration (Figure 4B). The confusion matrix showed correct classification for all AC and LC samples, whereas only one TC sample was assigned to AC (Figure 4C). This limited misclassification was also evident in the sample-wise classification quality plot (Figure 4D).

**Figure 4:**
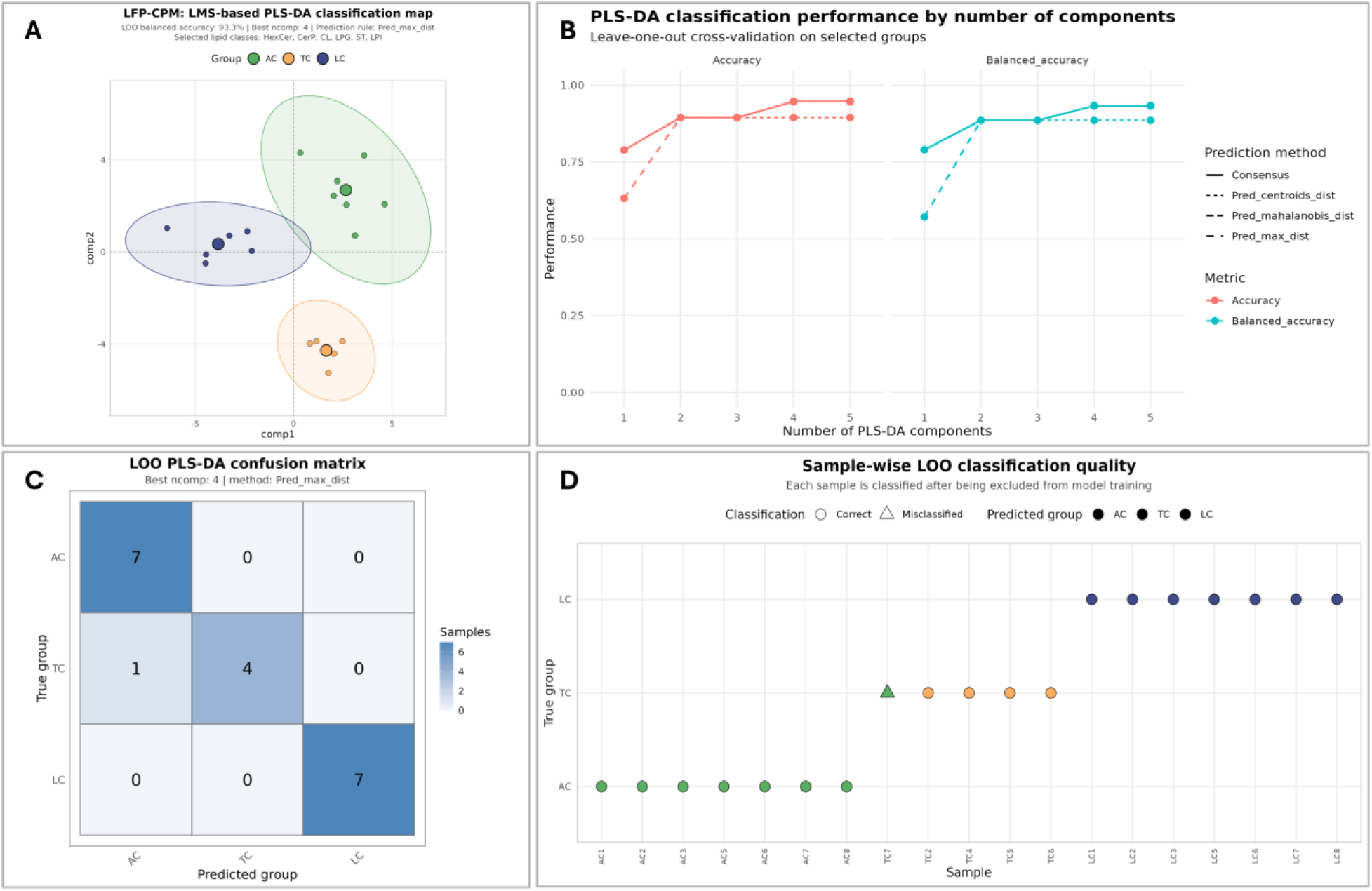
LFP-based geographical-origin classification of camel milk samples using the selected lipid-class subset. (A) PLS-DA classification map showing the separation of the three authenticated regional groups: AC, TC and LC. (B) Cross-validation performance across different numbers of PLS-DA components, reported as accuracy and balanced accuracy. (C) Confusion matrix for the selected model. (D) Sample-wise classification quality, showing correctly classified and misclassified samples.

The comparison with the all-class analysis further highlighted the usefulness of class-aware feature selection. Although the model based on all lipid classes still retained classification ability, it showed broader group dispersion and greater overlap among regional profiles. By contrast, the selected-class model provided a cleaner separation of the three geographical origins. This suggests that class-aware in silico hydrolysis can improve origin-oriented workflows not simply by reducing the number of variables, but by allowing users to remove lipid classes that dilute or obscure the most relevant food-origin signal.

The Unknown Sample Classifier was then used to simulate an authenticity-oriented scenario. A subset of samples was treated as unknown and projected into the reference LFP index space. The classifier assigned AC4 to AC, TC1 and TC8 to TC, and LC4 to LC (Figure 5B–C). Most assignments showed high confidence and full consensus, whereas TC1 displayed lower confidence and moderate reliability, suggesting a less clearly defined position relative to the reference groups.

**Figure 5:**
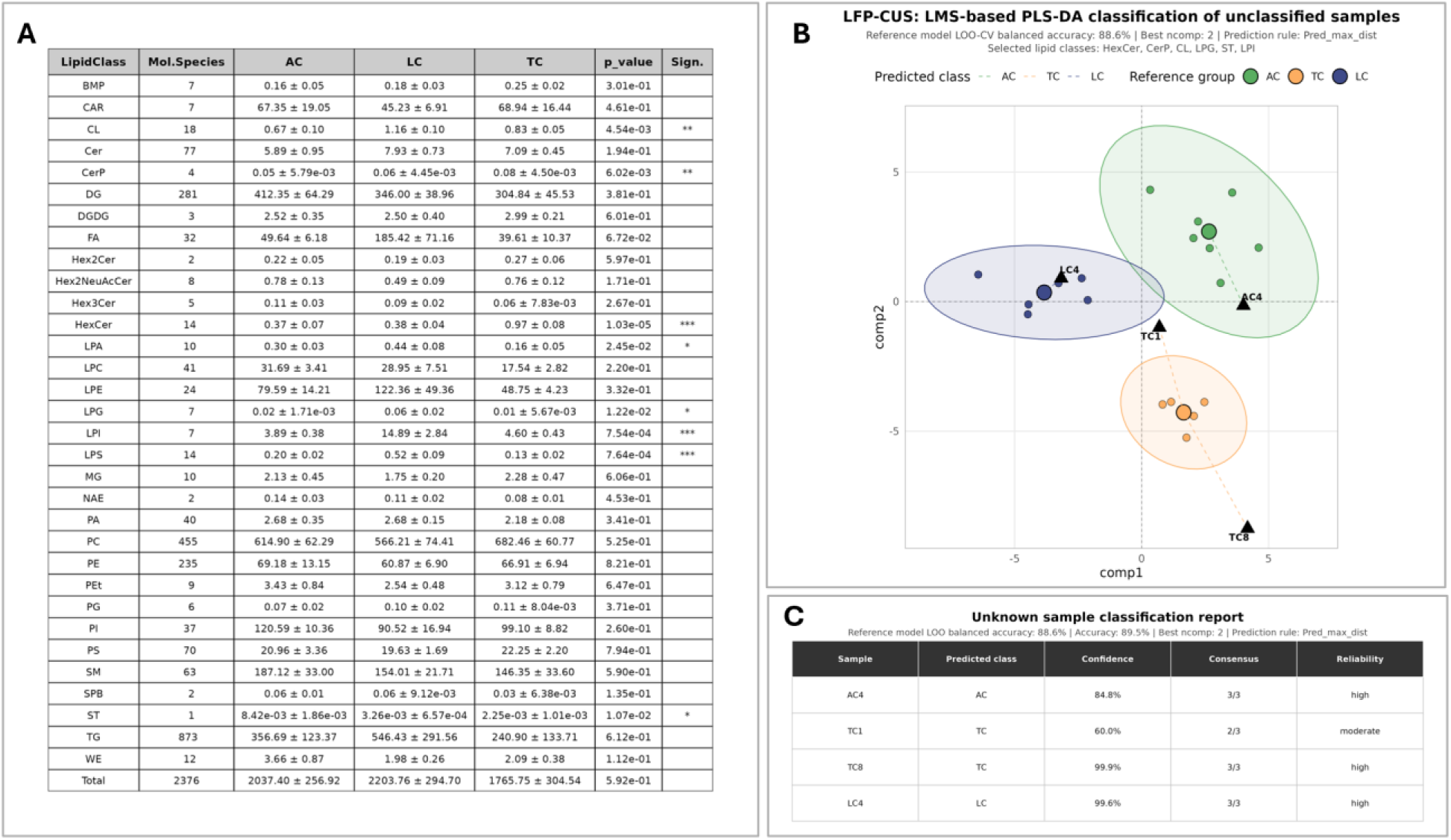
Lipid-class selection and unknown-sample assignment in the camel milk case study. (A) Lipid-class summary across the three geographical origins, used to identify the lipid classes most informative for regional discrimination. (B) LFP-based projection of unknown samples into the authenticated reference space. (C) Unknown-sample classification report showing predicted class, confidence, consensus and reliability for each unclassified sample.

Overall, the camel milk case study demonstrates that LFP can recover the regional structure described in the original lipidomics study while adding two complementary advantages: first, the transformation of molecular lipidomics data into interpretable food lipid indices; second, the possibility of using class-aware in silico hydrolysis to focus the analysis on lipid classes that are most informative for geographical-origin assessment. This remains an exploratory application and requires larger independent validation cohorts before routine authenticity use, but it illustrates how LFP can support traceability-oriented food lipidomics in an interpretable way.

## 5. Discussion: added value and limitations of LFP

The three case studies presented in this work illustrate the complementary value of LFP for the interpretation of LC/MS-based food lipidomics datasets. In all cases, the LFP analysis recovered the main conclusions of the original studies but reorganized them into a smaller number of food-oriented interpretative domains. This is an important point because LFP is not intended to generate an alternative or competing interpretation of lipidomics data. Rather, it provides an additional layer that translates molecular-species information into indices more directly connected to food lipid composition, oxidative stability, processing effects and traceability.

In the *Chlorella vulgaris* irradiation dataset, the original study described a lipid remodelling response involving fatty acids, membrane lipids and storage lipids. LFP retained this interpretation, but expressed it through coordinated changes in chain remodelling, compositional balance and oxidative stability. In the bottarga dataset, the original spatial lipidomics analysis identified clear differences between the inner and outer layers of the product, consistent with differential preservation, hydrolysis and oxidative exposure during processing. LFP recovered this inner–outer distinction and added a compact interpretation based on chain architecture, compositional balance and oxidative susceptibility. In the camel milk dataset, the original work reported regional lipid variation across three geographical origins. LFP preserved this geographical information and showed that it could be used in an exploratory origin-assessment workflow, while linking sample classification to interpretable lipid indices.

A distinctive advantage of LFP is that in silico hydrolysis is class-aware. Reconstructed chains are not collapsed into an anonymous fatty-acid pool but remain linked to the lipid classes from which they derive. This allows the lipidome to be explored at different levels of resolution. Indices may first be calculated across the complete set of lipid classes to provide an overall description of lipid composition. When a more specific foodomics question is addressed, the analysis can instead be restricted to selected lipid classes, excluding those that are poorly informative or unrelated to the question under investigation. The camel milk case study illustrates this flexibility: restricting the analysis to lipid classes showing regional differences produced a clearer and more stable classification space than the all-class analysis, while preserving the interpretability of the resulting index profile.

This aspect differentiates LFP from conventional fatty-acid profiling. Classical GC-based fatty-acid analysis remains essential for standardized and quantitative fatty-acid determination, especially when regulatory or nutritional labelling purposes are involved. However, chemical hydrolysis removes the molecular and lipid-class context of the original lipidome. LFP does not replace this approach but complements it by exploiting the structural information already present in annotated LC/MS lipidomics datasets. In this way, it enables food lipidomes to be interpreted not only in terms of individual lipid species, but also through lipid-quality, omega-balance, oxidative-stability, chain-remodelling and ether-linked-chain indices.

The proposed workflow may be particularly useful in studies where the aim is not only to identify discriminant molecules, but also to understand the type of lipid change occurring across samples, treatments, product layers or geographical origins. For this reason, LFP may facilitate communication between lipidomics specialists and food scientists, providing a common interpretative language that connects high-resolution lipidomics data with concepts commonly used in food chemistry and food quality assessment.

Some limitations should also be considered. LFP depends on the quality and structural resolution of lipid annotations: datasets containing only lipid-class abundances or unresolved sum compositions are not sufficient for complete chain reconstruction. In addition, LC/MS lipidomics data are often semi-quantitative and may be affected by lipid-class-dependent ionization efficiency, extraction bias and incomplete lipidome coverage. Therefore, LFP-derived indices should be interpreted as indices of the analysed lipidomics dataset rather than as absolute measures of food composition. Finally, classification-oriented applications, such as geographical-origin or authenticity assessment, require larger independent validation cohorts before they can be used in routine quality-control contexts.

## 6. Conclusions

Lipid Food Profile expands the LipidOne platform by providing an index-based strategy for the interpretation of LC/MS food lipidomics datasets. Through class-aware in silico hydrolysis, annotated intact lipid species are translated into reconstructed chain-level information while preserving lipid-class origin. This allows complex lipidomics datasets to be summarized into food-relevant indices related to lipid quality, compositional balance, omega balance, oxidative stability, chain remodelling and ether-linked chain contribution.

Across the three case studies, LFP recovered the main findings of the original lipidomics studies and reorganized them into compact, chemically meaningful food lipid profiles. In *Chlorella vulgaris*, the module summarized irradiation-associated lipid remodelling; in *Mugil cephalus* bottarga, it captured spatial lipid heterogeneity between inner and outer layers; and in camel milk, it supported an exploratory geographical-origin assessment based on interpretable lipid indices.

LFP is not intended to replace conventional fatty-acid analysis, targeted lipid quantification or external validation of authenticity markers. Rather, it provides a complementary interpretative layer for intact-lipid LC/MS datasets, helping users move from long molecular-species tables to structured lipid profiles that are easier to compare across foods, treatments, processing conditions and geographical origins.

## Acknowledgements

We would like to thank Matteo Boschi (UMMON.it) for his invaluable cooperation in developing the LipidOne 2.5 website front-end and coding its back-end interactions with the R scripts.

## Author contributions

Conceptualisation: RMP, DFM and HBRA. Software development: RMP. Data analysis: DFM. Data validation: DFM and HBRA. Data interpretation: RMP, DFM and HBRA. Writing—original draft: DFM and RMP. Writing—review and editing: RMP, DFM and HBRA. All authors read and approved the final manuscript.

## Data availability

The original lipidomics datasets analysed in this study are publicly available through LipidOne platform as example dataset.

