## supplementary material for "From molecular lipidomics to interpretable food lipid profiles: the Lipid Food Profile module in LipidOne"

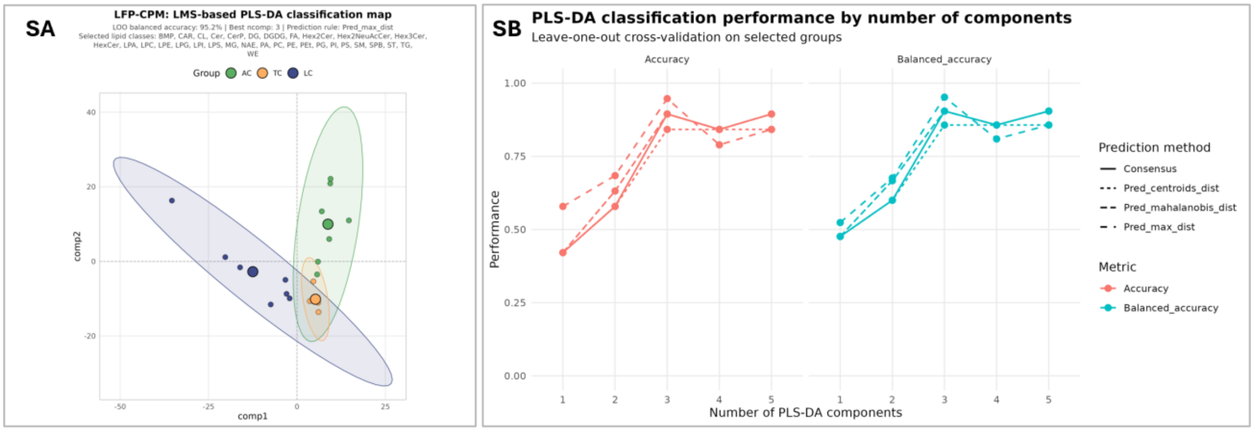


**Figure S1: PLS-DA classification of camel milk geographical origin using all lipid classes**. (A) PLS-DA classification map obtained using the complete lipid-class set detected in the dataset, including lipid classes not reaching statistical significance in the initial regional comparison. (B) Cross-validation performance across different numbers of PLS-DA components, reported as accuracy and balanced accuracy for the available prediction methods. Although the all-class model showed high classification performance, the score map displayed broader group dispersion and partial overlap among regional profiles compared with the significant-class subset used in the main analysis. AC, Fuhai County; TC, Yumin County; LC, Huocheng County.
