## Supplementary Table S1 for "From molecular lipidomics to interpretable food lipid profiles: the Lipid Food Profile module in LipidOne"

**Lipid Food Profile indices, formulas, interpretive rules, biological rationale, and supporting references.**

This table reports the complete set of indices implemented in the LFP module, including their category assignment, index code, display name, formula, direction-specific interpretive phrases, biological explanation, and supporting references.

| Category | Index Code | Index Name | Formula | Interpretive phrase if increased | Interpretive phrase if decreased | Biological explanation | References |
|---|---|---|---|---|---|---|---|
| Food Lipid Quality | AI | Atherogenic Index (AI) | [C12:0 + (4 × C14:0) + C16:0] / [ΣMUFA + ΣPUFA] | Higher atherogenic balance in EXP suggests a less favourable saturated to unsaturated chain profile. | Lower atherogenic balance in EXP suggests a more favourable saturated to unsaturated chain profile. | The Atherogenic Index (AI) is a fatty-acid-based lipid-quality index that describes the relative contribution of the main atherogenic saturated fatty acids — lauric acid (C12:0), myristic acid (C14:0), and palmitic acid (C16:0) — in relation to the total unsaturated fatty-acid fraction of a food. Within the Lipid Food Profile framework, AI should be interpreted as a compositional index of the food lipid profile rather than as a direct clinical risk marker. A higher AI indicates that the lipid fraction is relatively more enriched in selected saturated fatty acids and less supported by MUFA and PUFA, suggesting a less favourable lipid-quality profile. Conversely, a lower AI reflects a greater contribution of unsaturated fatty acids and indicates a more favourable fatty-acid balance. This index is particularly useful for comparing food matrices, raw versus processed/cooked products, different formulations, animal feeding systems, or technological conditions that may modify the saturated-to-unsaturated fatty-acid balance. | (J. Chen & Liu, 2020; Dal Bosco et al., 2024) |
| | TI | Thrombogenicity Index (TI) | [C14:0 + C16:0 + C18:0] / [(0.5 × ΣMUFA) + (0.5 × Σn-6 PUFA) + (3 × Σn-3PUFA) + (Σn-3 PUFA / Σn-6 PUFA)] | Higher thrombogenic balance in EXP indicates a less favourable overall fatty acid quality pattern. | Lower thrombogenic balance in EXP indicates a more favourable overall fatty acid quality pattern. | The Thrombogenicity Index (TI) is a fatty-acid-based lipid-quality index that describes the balance between selected saturated fatty acids associated with a more thrombogenic profile — myristic acid (C14:0), palmitic acid (C16:0), and stearic acid (C18:0) — and a weighted pool of unsaturated fatty acids considered more protective, including MUFA, n-6 PUFA, n-3 PUFA, and the n-3/n-6 PUFA ratio. Within the Food Lipid Profile framework, TI should be interpreted as a compositional index of the food lipid profile rather than as a direct clinical risk marker. A higher TI indicates that the food lipid fraction is relatively more enriched in saturated fatty acids associated with thrombogenic potential and less supported by protective unsaturated fatty acids, suggesting a less favourable lipid-quality profile. Conversely, a lower TI reflects a greater contribution of unsaturated fatty acids, particularly n-3 PUFA, and indicates a more favourable fatty-acid balance. This index is useful for comparing food matrices, processing conditions, raw versus cooked products, animal feeding systems, or reformulated foods that may differ in their saturated, monounsaturated, and polyunsaturated fatty-acid composition. | (Dal Bosco et al., 2024; Paszczyk & Czarnowska-Kujawska, 2022; Rashidimehr et al., 2025) |
| | h_H | Hypo-/Hyper-cholesterolemic Ratio | [C18:1 + ΣPUFA] / [C12:0 + C14:0 + C16:0] | Higher h/H ratio in EXP indicates enrichment of hypocholesterolemic relative to hypercholesterolemic chains. | Lower h/H ratio in EXP indicates a less favorable hypocholesterolemic to hypercholesterolemic chain balance. | The H/H ratio is a fatty-acid-based lipid-quality index that describes the balance between unsaturated fatty acids, represented by cis-C18:1 and total PUFA, and selected saturated fatty acids, represented by C12:0, C14:0, and C16:0. Within the Food Lipid Profile framework, this index should be interpreted as a compositional index of the food lipid profile. A higher H/H ratio indicates that the lipid fraction is relatively more enriched in oleic acid and polyunsaturated fatty acids, suggesting a more favourable unsaturated-fatty-acid contribution to the overall lipid quality of the food. Conversely, a lower H/H ratio reflects a greater relative contribution of selected saturated fatty acids and indicates a less favourable fatty-acid balance. This index is useful for comparing food matrices, processing conditions, formulations, animal feeding systems, or other factors that may modify the relationship between unsaturated and saturated fatty acids in foods. | (J. Chen & Liu, 2020; Santos-Silva et al., 2002) |
| Chain Remodelling | C16_DI | C16 Desaturation Index | C16:1 / (C16:0 + C16:1) | Higher C16 desaturation in EXP suggests stronger conversion toward monounsaturated C16 chains. | Lower C16 desaturation in EXP suggests weaker conversion toward monounsaturated C16 chains. | The C16 Desaturation Index (C16_DI) is a chain-specific fatty-acid remodelling index that expresses the relative contribution of monounsaturated C16 fatty acids to the total C16 fatty-acid pool. It is calculated as C16:1/(C16:0 + C16:1), where C16:0 represents palmitic acid and C16:1 represents the corresponding monounsaturated C16 fraction, mainly palmitoleic acid when positional isomers are resolved. Within the Food Lipid Profile framework, C16_DI describes whether the C16 lipid fraction is more strongly characterized by saturated or monounsaturated C16 chains. A higher C16_DI indicates a greater relative contribution of C16:1 and suggests a shift of the C16 pool toward monounsaturated chains. Conversely, a lower C16_DI reflects a stronger predominance of C16:0 and a lower relative contribution of monounsaturated C16 species. This index is useful for comparing food matrices, animal genotypes, feeding systems, formulations, or processing conditions that may modify the balance between palmitic and palmitoleic acid-related lipid fractions. | (Inostroza et al., 2022) |
| | C18_DI | C18 Desaturation Index | C18:1 / (C18:0 + C18:1) | Higher C18 desaturation in EXP suggests stronger conversion toward monounsaturated C18 chains. | Lower C18 desaturation in EXP suggests weaker conversion toward monounsaturated C18 chains. | The C18 Desaturation Index (C18_DI) is a chain-specific remodelling index that expresses the relative contribution of monounsaturated C18 fatty acids to the combined C18:0 and C18:1 pool. It is calculated as C18:1/(C18:0 + C18:1), where C18:0 represents stearic acid and C18:1 represents the monounsaturated C18 fraction, mainly oleic acid when positional isomers are resolved. Within the Food Lipid Profile framework, C18_DI describes whether the C18 lipid fraction is more strongly characterized by saturated or monounsaturated C18 chains. A higher C18_DI indicates a stronger contribution of C18:1 and suggests a C18 profile shifted toward monounsaturated chains, whereas a lower C18_DI reflects a greater predominance of C18:0 and a lower relative contribution of monounsaturated C18 species. This index is useful for comparing food matrices, animal genotypes, feeding systems, formulations, processing conditions, or other factors that may modify the balance between stearic-acid- and oleic-acid-related lipid fractions. | (Inostroza et al., 2022) |

| | | | | | | | |
|---|---|---|---|---|---|---|---|
| Omega Balance | DHA_ARA | DHA/ARA Ratio | C22:6 n-3 / C20:4 n-6 | Higher DHA/ARA ratio in EXP indicates a more omega 3 oriented long chain PUFA balance. | Lower DHA/ARA ratio in EXP indicates a weaker omega 3 contribution relative to arachidonic acid. | The DHA/ARA ratio is a targeted long-chain PUFA balance index that compares docosahexaenoic acid (DHA; C22:6n-3) with arachidonic acid (ARA; C20:4n-6). Within the Food Lipid Profile framework, this index describes whether the long-chain PUFA fraction of a food is more oriented toward an omega-3 DHA-related profile or toward an omega-6 ARA-related profile. A higher DHA/ARA ratio indicates a greater relative contribution of DHA compared with ARA, suggesting a long-chain PUFA pattern more strongly characterized by DHA. Conversely, a lower DHA/ARA ratio reflects a weaker DHA contribution relative to arachidonic acid and indicates a long-chain PUFA profile more influenced by the ARA component. | (Araújo et al., 2022) |
| | EPA_ARA | EPA/ARA Ratio | C20:5 n-3 / C20:4 n-6 | Higher EPA/ARA ratio in EXP indicates a more omega 3 oriented long chain PUFA balance. | Lower EPA/ARA ratio in EXP indicates a weaker omega 3 contribution relative to arachidonic acid. | The EPA/ARA ratio is a targeted long-chain PUFA balance index that compares eicosapentaenoic acid (EPA; C20:5n-3) with arachidonic acid (ARA; C20:4n-6). Within the Food Lipid Profile framework, this index describes whether the long-chain PUFA fraction of a food is more oriented toward an omega-3 EPA-related profile or toward an omega-6 ARA-related profile. A higher EPA/ARA ratio indicates a greater relative contribution of EPA compared with ARA, suggesting a long-chain PUFA pattern more strongly characterized by EPA. Conversely, a lower EPA/ARA ratio reflects a weaker EPA contribution relative to arachidonic acid and indicates a long-chain PUFA profile more influenced by the ARA component. | (Araújo et al., 2022) |
| | EPA+DHA Fraction | EPA+DHA Fraction | (EPA + DHA) / total reconstructed chain signal | Higher EPA+DHA Fraction in EXP reflects greater EPA and DHA contribution to the reconstructed chain pool. | Lower EPA+DHA Fraction in EXP reflects reduced EPA and DHA contribution to the reconstructed chain pool. | The EPA+DHA Fraction is a lipidome-derived compositional index that estimates the relative contribution of EPA and DHA to the total reconstructed fatty-acyl chain signal of a food lipid profile. It is calculated as (EPA + DHA)/total reconstructed chain signal, where EPA and DHA represent the main long-chain omega-3 PUFA typically used to characterize marine and omega-3-rich food matrices. Within the Food Lipid Profile framework, a higher EPA+DHA Fraction indicates that EPA- and DHA-containing chains contribute more strongly to the reconstructed lipidome, suggesting a lipid profile more enriched in long-chain omega-3 PUFA. Conversely, a lower value reflects a reduced EPA+DHA contribution and indicates that other fatty-acyl chains dominate the reconstructed lipid pool. This index is particularly useful for comparing fish, seafood, marine oils, omega-3-enriched products, aquaculture-derived foods, and formulations or feeding systems that modify EPA and DHA content. | (J. Chen & Liu, 2020; Nava et al., 2023; Sprague et al., 2016) |
| Compositional Balance | LCFA_pct | Long-Chain FA % | 100 × Σlong-chain fatty-acid chains / total reconstructed chain signal | Higher long chain fatty acid fraction in EXP indicates a shift toward longer acyl chain composition. | Lower long chain fatty acid fraction in EXP indicates a reduced contribution of long acyl chains. | The Long-Chain Fatty Acid Percentage (LCFA_pct) is a compositional balance index that estimates the relative contribution of long-chain fatty-acid residues to the total reconstructed fatty-acyl chain signal of a food lipid profile. It is calculated as 100 × Σlong-chain fatty-acid chains / total reconstructed chain signal. Within the Food Lipid Profile framework, LCFA_pct describes whether the reconstructed lipidome is more strongly represented by long acyl chains or by shorter-chain components. A higher LCFA_pct indicates a greater contribution of long-chain fatty-acid residues and suggests a shift toward longer acyl-chain composition. Conversely, a lower LCFA_pct reflects a reduced contribution of long-chain residues and may indicate a relatively greater contribution of short- or medium-chain fatty acids, depending on the food matrix and the fatty acids included in the denominator. | (Paszczyk et al., 2019) |
| | MCFA_LCFA | MCFA/LCFA Ratio | Σmedium-chain fatty-acid chains / Σlong-chain fatty-acid chains | Higher MCFA/LCFA ratio in EXP indicates a greater medium chain contribution relative to long chains. | Lower MCFA/LCFA ratio in EXP indicates a lower medium chain contribution relative to long chains. | The MCFA/LCFA ratio is a chain-length compositional index that compares the relative contribution of medium-chain fatty-acid residues with that of long-chain fatty-acid residues in a food lipid profile. It is calculated as Σmedium-chain fatty-acid chains / Σlong-chain fatty-acid chains. Within the Food Lipid Profile framework, this index describes whether the reconstructed lipidome is more strongly characterized by medium-chain or long-chain acyl residues. A higher MCFA/LCFA ratio indicates a greater medium-chain contribution relative to long-chain residues, suggesting a lipid profile more enriched in medium-chain components. Conversely, a lower MCFA/LCFA ratio reflects a lower medium-chain contribution and a stronger predominance of long-chain lipid species. This index is useful for distinguishing food matrices enriched in medium-chain lipids from those dominated by longer-chain acyl residues, but its interpretation depends on the food matrix and on the chain-length cut-offs used to define MCFA and LCFA. | (Sun et al., 2013) |
| | MCFA_pct | Medium-Chain FA % | 100 × Σmedium-chain fatty-acid chains / total reconstructed chain signal | Higher medium chain fatty acid fraction in EXP indicates enrichment of shorter acyl chains. | Lower medium chain fatty acid fraction in EXP indicates depletion of shorter acyl chains. | The Medium-Chain Fatty Acid Percentage (MCFA_pct) is a chain-length compositional index that estimates the relative contribution of medium-chain fatty-acid residues to the total reconstructed fatty-acyl chain signal of a food lipid profile. It is calculated as 100 × Σmedium-chain fatty-acid chains / total reconstructed chain signal. Within the Food Lipid Profile framework, MCFA_pct describes whether the reconstructed lipidome is enriched in medium-length acyl residues. A higher MCFA_pct indicates a greater contribution of medium-chain fatty acids and suggests a lipid profile more strongly characterized by medium-chain components. Conversely, a lower MCFA_pct reflects a reduced contribution of medium-chain residues and indicates that other chain-length groups contribute more strongly to the reconstructed lipid pool. This index can be particularly relevant for dairy-derived matrices, coconut-derived products, palm-kernel-derived fats, structured lipids, and formulated foods naturally or technologically enriched in medium-chain lipids. | (Paszczyk et al., 2019) |
| | MUFA_SFA | MUFA/SFA Ratio | Σmonounsaturated fatty-acid chains / Σsaturated fatty-acid chains | Higher MUFA/SFA ratio in EXP suggests a more favourable monounsaturated to saturated chain balance. | Lower MUFA/SFA ratio in EXP suggests a less favourable monounsaturated to saturated chain balance. | The MUFA/SFA ratio is a compositional balance index that compares the relative contribution of monounsaturated fatty-acid residues with that of saturated fatty-acid residues in a food lipid profile. It is calculated as ΣMUFA/ΣSFA, where MUFA are fatty-acid chains containing one double bond and SFA are fatty-acid chains without double bonds. Within the Food Lipid Profile framework, this index describes whether the lipid profile is more strongly characterized by monounsaturated or saturated chain composition. A higher MUFA/SFA ratio indicates a greater monounsaturated contribution relative to saturated fatty acids, suggesting a lipid profile more enriched in monoenoic chains. Conversely, a lower MUFA/SFA ratio reflects a stronger saturated-chain contribution and a lower relative presence of monounsaturated residues. | (Yazdanparast et al., 2025) |
| | PUFA_MUFA | PUFA/MUFA Ratio | Σpolyunsaturated fatty-acid chains / Σmonounsaturated fatty-acid chains | Higher PUFA/MUFA ratio in EXP indicates greater polyunsaturated contribution relative to monounsaturated chains. | Lower PUFA/MUFA ratio in EXP indicates reduced polyunsaturated contribution relative to monounsaturated chains. | The PUFA/SFA ratio is a fatty-acid-based lipid-quality index that describes the balance between total polyunsaturated fatty acids and total saturated fatty acids in a food matrix. Within the Food Lipid Profile framework, this index should be interpreted as a broad compositional index of the food lipid profile. A higher PUFA/SFA ratio indicates that the lipid fraction is relatively more enriched in polyunsaturated fatty acids, suggesting a fatty-acid profile more strongly characterized by unsaturation. Conversely, a lower PUFA/SFA ratio reflects a greater relative contribution of saturated fatty acids and indicates a lipid profile more dominated by saturation. This index is useful for comparing food matrices, raw versus cooked products, formulations, processing conditions, animal feeding systems, or other factors that may modify the relative abundance of PUFA and SFA in foods. | (Hamdy et al., 2026) |

| | | | | | | | |
|---|---|---|---|---|---|---|---|
| | PUFA_SFA | PUFA/SFA Ratio | Σpolyunsaturated fatty-acid chains / Σsaturated fatty-acid chains | Higher PUFA/SFA ratio in EXP suggests a more favourable polyunsaturated to saturated chain balance. | Lower PUFA/SFA ratio in EXP suggests a less favourable polyunsaturated to saturated chain balance. | The PUFA/SFA ratio is a fatty-acid compositional balance index that compares the relative contribution of polyunsaturated fatty-acid residues with that of saturated fatty-acid residues in a food lipid profile. It is calculated as ΣPUFA/ΣSFA, where PUFA are fatty-acid chains containing two or more double bonds and SFA are fatty-acid chains without double bonds. Within the Food Lipid Profile framework, this index describes whether the lipid profile is more strongly characterized by polyunsaturated or saturated chain composition. A higher PUFA/SFA ratio indicates a greater polyunsaturated contribution relative to saturated fatty acids, suggesting a lipid profile more enriched in polyunsaturated chains. Conversely, a lower PUFA/SFA ratio reflects a stronger saturated-chain contribution and a lower relative presence of polyunsaturated residues. This index is useful for comparing food matrices, formulations, processing conditions, or production systems that modify the balance between PUFA and SFA. | (J. Chen & Liu, 2020) |
| | SFA_pct | Saturated FA % | 100 × Σsaturated fatty-acid chains / total reconstructed chain signal | Higher saturated fatty acid fraction in EXP indicates a shift toward a more saturated lipid matrix. | Lower saturated fatty acid fraction in EXP indicates a shift away from saturated chains. | The Saturated Fatty Acid Percentage (SFA_pct) is a class-level compositional index that estimates the relative contribution of saturated fatty-acid residues to the total reconstructed fatty-acyl chain signal of a food lipid profile. It is calculated as 100 × Σsaturated fatty-acid chains / total reconstructed chain signal. Within the Food Lipid Profile framework, SFA_pct describes whether the reconstructed lipidome is more strongly characterized by saturated chains. A higher SFA_pct indicates a greater saturated-chain contribution and suggests a lipid matrix more enriched in saturated fatty-acid residues. Conversely, a lower SFA_pct reflects a reduced saturated-chain contribution and indicates a shift away from saturated residues toward other fatty-acid classes. | (Kaçar et al., 2024) |
| Oxidative Stability | PI | Peroxidability Index | Σ[relative chain abundance × peroxidability coefficient by number of double bonds] | Higher peroxidability index in EXP suggests greater susceptibility to oxidative lipid damage. | Lower peroxidability index in EXP suggests reduced susceptibility to oxidative lipid damage. | The Peroxidability Index (PI) is an oxidative stability index that estimates the theoretical susceptibility of a food lipid profile to lipid peroxidation. It is calculated by summing the relative abundance of fatty-acid chains weighted by peroxidability coefficients based on the number of double bonds. Within the Food Lipid Profile framework, PI describes the intrinsic oxidative sensitivity of the reconstructed lipidome: lipid profiles enriched in highly unsaturated fatty acids, especially long-chain PUFA such as EPA and DHA, generally show higher PI values, whereas profiles dominated by saturated, monounsaturated, or less unsaturated chains show lower values. A higher PI indicates a lipid matrix with greater theoretical susceptibility to oxidative lipid damage, while a lower PI indicates reduced compositional susceptibility to peroxidation. This index is particularly useful for evaluating food matrices rich in PUFA, such as fish, seafood, marine oils, omega-3-enriched products, nuts, seeds, and processed foods where oxidative stability is a key quality dimension. | (Yun & Surh, 2012) |
| | UI | Unsaturation Index | Σ[relative chain abundance × number of double bonds] | Higher unsaturation index in EXP indicates a more unsaturated chain profile. | Lower unsaturation index in EXP indicates a more saturated chain profile. | The Unsaturation Index (UI) is a weighted structural index of the fatty-acid profile that estimates the overall degree of unsaturation of a food lipid fraction. It is calculated by multiplying the relative abundance of each unsaturated fatty acid by its number of double bonds and summing these weighted contributions across the lipid profile. A higher UI indicates that the food lipid profile is more enriched in highly unsaturated fatty acids, such as long-chain polyunsaturated species, whereas a lower UI reflects a lipid profile with a lower contribution of highly unsaturated fatty acids and a stronger relative presence of saturated, monounsaturated, or less unsaturated species. This index is useful for comparing food matrices, raw versus cooked products, processing conditions, formulations, storage conditions, or production systems that may modify the structural unsaturation of food lipids. | (J. Chen & Liu, 2020) |
| Ether-Linked Chains | Ether_fraction | Ether Chain Fraction | Σether-linked chains (O- + P-) / total reconstructed chain signal | Higher ether linked chain fraction in EXP suggests enrichment of ether lipid derived chains. | Lower ether linked chain fraction in EXP suggests reduced contribution of ether lipid derived chains. | The Ether Chain Fraction is an ether-linked lipid index that estimates the relative contribution of ether-derived chains to the total reconstructed chain signal of a food lipid profile. It is calculated as Σether-linked chains / total reconstructed chain signal, where O- denotes alkyl ether-linked chains and P- denotes vinyl ether/plasmalogen-related chains according to the lipid annotation rules used by the module. Within the Food Lipid Profile framework, this index describes whether the reconstructed lipidome is enriched in ether-linked lipid species rather than conventional ester-linked acyl chains. A higher Ether Chain Fraction indicates a greater contribution of ether-linked species and suggests enrichment of ether lipid-derived chains, whereas a lower value reflects reduced contribution of ether-linked lipid species. This index is particularly useful for comparing animal-derived foods, seafood, organ-rich matrices, and lipid profiles where ether phospholipids or plasmalogens are relevant components of the food lipidome. | (Z. Chen et al., 2025) |
| | Alkyl_fraction | Alkyl Chain Fraction | Σalkyl ether-linked chains (O-) / total reconstructed chain signal | Higher alkyl ether fraction in EXP indicates enrichment of plasmanyl related chains. | Lower alkyl ether fraction in EXP indicates reduced contribution of plasmanyl related chains. | The Alkyl Chain Fraction is an ether-linked chain index that estimates the relative contribution of alkyl ether-linked chains to the total reconstructed chain signal of a food lipid profile. It is calculated as Σalkyl ether-linked chains/ total reconstructed chain signal, where O- denotes plasmanyl-type alkyl ether-linked species according to the lipid annotation rules. Within the Food Lipid Profile framework, this index describes whether the reconstructed lipidome is enriched in O-linked ether lipid-derived chains. A higher Alkyl Chain Fraction indicates a greater contribution of plasmanyl-related chains, whereas a lower value reflects reduced alkyl ether-linked chain representation. | (Z. Chen et al., 2025) |
| | Alkenyl_fraction | Alkenyl Chain Fraction | Σalkenyl/vinyl-ether-linked chains (P-) / total reconstructed chain signal | Higher alkenyl ether fraction in EXP indicates enrichment of plasmalogen related chains. | Lower alkenyl ether fraction in EXP indicates reduced contribution of plasmalogen related chains. | The Alkenyl Chain Fraction is an ether-linked chain index that estimates the relative contribution of alkenyl/vinyl-ether-linked chains to the total reconstructed chain signal of a food lipid profile. It is calculated as Σalkenyl/vinyl-ether-linked chains (P-) / total reconstructed chain signal, where P- denotes plasmenyl or plasmalogen-related species according to the lipid annotation rules used by the module. Within the Food Lipid Profile framework, this index describes whether the reconstructed lipidome is enriched in plasmalogen-derived chains. A higher Alkenyl Chain Fraction indicates a greater contribution of P-linked plasmalogen-related chains, whereas a lower value reflects reduced plasmalogen-related chain representation. | (Z. Chen et al., 2025) |

**Reference Table S1**


Araújo, B. C., Skrzynska, A. K., Marques, V. H., Tinajero, A., Del Rio-Zaragoza, O. B., Viana, M. T., & Mata-Sotres, J. A. (2022). Dietary Arachidonic Acid (20:4n-6) Levels and Its Effect on Growth Performance, Fatty Acid Profile, Gene Expression for Lipid Metabolism, and Health Status of Juvenile California Yellowtail (Seriola dorsalis). *Fishes*, *7*(4), 185. https://doi.org/10.3390/fishes7040185

Chen, J., & Liu, H. (2020). Nutritional Indices for Assessing Fatty Acids: A Mini-Review. *International Journal of Molecular Sciences*, *21*(16), 5695. https://doi.org/10.3390/ijms21165695

Chen, Z., Dong, C., Chen, L., Song, M., Zhou, X., Lv, D., & Li, Q. (2025). The Changes in Plasmalogens: Chemical Diversity and Nutritional Implications—A Narrative Review. *Nutrients*, *17*(22), 3497. https://doi.org/10.3390/nu17223497

Dal Bosco, A., Cavallo, M., Menchetti, L., Angelucci, E., Cartoni Mancinelli, A., Vaudo, G., Marconi, S., Camilli, E., Galli, F., Castellini, C., & Mattioli, S. (2024). The Healthy Fatty Index Allows for Deeper Insights into the Lipid Composition of Foods of Animal Origin When Compared with the Atherogenic and Thrombogenicity Indexes. *Foods*, *13*(10), 1568. https://doi.org/10.3390/foods13101568

Hamdy, S. M., Metwally, A. A., Deghedi, M. A., & Abdelmontaleb, H. S. (2026). Comparative effects of natural and functional lipid sources on physicochemical, textural, sensory, and nutritional lipid indices of spreadable processed cheese analogue. *Scientific Reports*, *16*(1), 17326. https://doi.org/10.1038/s41598-026-55188-3

Inostroza, K., Larama, G., Bravo, S., Díaz, M., & Sepúlveda, N. (2022). Utilization of Wool Integral Lipids to Determine Milk Fat Content in Suffolk Down Ewes. *Applied Sciences*, *12*(3), 1046. https://doi.org/10.3390/app12031046

Kaçar, S., Kayhan Kaya, H., & Başhan, M. (2024). Seasonal Changes in Fatty Acid Composition of *Chondrostoma regium* Lipids. *Polish Journal of Food and Nutrition Sciences*, 387–398. https://doi.org/10.31883/pjfns/195435

Nava, V., Turco, V. L., Licata, P., Panayotova, V., Peycheva, K., Fazio, F., Rando, R., Di Bella, G., & Potortì, A. G. (2023). Determination of Fatty Acid Profile in Processed Fish and Shellfish Foods. *Foods*, *12*(13), 2631. https://doi.org/10.3390/foods12132631

Paszczyk, B., & Czarnowska-Kujawska, M. (2022). Fatty Acid Profile, Conjugated Linoleic Acid Content, and Lipid Quality Indices in Selected Yogurts Available on the Polish Market. *Animals*, *12*(1), 96. https://doi.org/10.3390/ani12010096

Paszczyk, B., Polak-Śliwińska, M., & Łuczyńska, J. (2019). Fatty Acids Profile, Trans Isomers, and Lipid Quality Indices in Smoked and Unsmoked Cheeses and Cheese-Like Products. *International Journal of Environmental Research and Public Health*, *17*(1), 71. https://doi.org/10.3390/ijerph17010071

Rashidimehr, A., Mohammadi-Nasrabadi, F., Alhouei, B., Khoshtinat, K., & Esfarjani, F. (2025). Fatty acid composition of dairy products and their impact on atherogenicity and thrombogenicity. *Scientific Reports*, *15*(1), 43613. https://doi.org/10.1038/s41598-025-27445-4

Santos-Silva, J., Bessa, R. J. B., & Santos-Silva, F. (2002). Effect of genotype, feeding system and slaughter weight on the quality of light lambs. *Livestock Production Science*, *77*(2–3), 187–194. https://doi.org/10.1016/S0301-6226(02)00059-3

Sprague, M., Dick, J. R., & Tocher, D. R. (2016). Impact of sustainable feeds on omega-3 long-chain fatty acid levels in farmed Atlantic salmon, 2006–2015. *Scientific Reports*, *6*(1), 21892. https://doi.org/10.1038/srep21892

Sun, Y., Bu, D. P., Wang, J. Q., Cui, H., Zhao, X. W., Xu, X. Y., Sun, P., & Zhou, L. Y. (2013). Supplementing different ratios of short- and medium-chain fatty acids to long-chain fatty acids in dairy cows: Changes of milk fat production and milk fatty acids composition. *Journal of Dairy Science*, *96*(4), 2366–2373. https://doi.org/10.3168/jds.2012-5356

Yazdanparast, S., Mohammadi-Nasrabadi, F., Hashmati, A., Rezazadeh, R., Taheri, M., Alhouei, B., & Esfarjani, F. (2025). Comprehensive assessment of fatty acid profiles of meat products to develop action plan strategies for healthier products. *Scientific Reports*, *15*(1), 23188. https://doi.org/10.1038/s41598-025-04749-z

Yun, J.-M., & Surh, J.-H. (2012). Fatty Acid Composition as a Predictor for the Oxidation Stability of Korean Vegetable Oils with or without Induced Oxidative Stress. *Preventive Nutrition and Food Science*, *17*(2), 158–165. https://doi.org/10.3746/pnf.2012.17.2.158
